# Inhibition of tRNA Synthetases Induces Persistence in *Chlamydia*

**DOI:** 10.1101/759084

**Authors:** Nathan D. Hatch, Scot P. Ouellette

**Author notes:** Corresponding Author: Department of Pathology and Microbiology, College of Medicine, University of Nebraska Medical Center, 985900 Nebraska Medical Center (DRC2 5022), Omaha, NE.

## Abstract

*Chlamydia trachomatis* is the leading cause of bacterial sexually transmitted infections, and *C. pneumoniae* causes community-acquired respiratory infections. *In vivo*, the host immune system will release interferon-gamma (IFNγ) to combat infection. IFNγ activates human cells to produce the tryptophan (trp) catabolizing enzyme, IDO. Consequently, there is a reduction in cytosolic trp in IFNγ-activated host cells. In evolving to obligate intracellular dependence, *Chlamydia* has significantly reduced its genome size and content as it relies on the host cell for various nutrients. Importantly, *C. trachomatis* and *C. pneumoniae* are trp auxotrophs and are starved for this essential nutrient when the human host cell is exposed to IFNγ. To survive this, chlamydiae enter an alternative growth state referred to as persistence. Chlamydial persistence is characterized by a halt in the division cycle, aberrant morphology, and, in the case of IFNγ-induced persistence, trp codon-dependent changes in transcription. We hypothesize that these changes in transcription are dependent on the particular amino acid starvation state. To investigate the chlamydial response mechanisms acting when other amino acids become limiting, we tested the efficacy of prokaryotic specific tRNA synthetase inhibitors, indolmycin and AN3365, to mimic starvation of trp and leucine, respectively. We show that these drugs block chlamydial growth and induce changes in morphology and transcription consistent with persistence. Importantly, growth inhibition was reversed when the compounds were removed from the medium. With these data, we find that indolmycin and AN3365 are valid tools that can be used to mimic the persistent state independently of IFNγ.

**Importance:** The obligate intracellular pathogen *Chlamydia trachomatis*, although treatable, remains a major public health concern due to rising infection rates. The asymptomatic nature of most *Chlamydia* infections is hypothesized to be a product of its ability to transition into a slow-growing state referred to as persistence. The most physiologically relevant inducer of persistence is the immune cytokine IFNγ, which in humans activates an enzyme that degrades tryptophan, an essential amino acid that *Chlamydia* scavenges from the host cell. Unfortunately, the exact timing at which *Chlamydia* is starved after IFNγ treatment is inexact. To mechanistically study persistence using genetic tools, an experimental model where amino acid starvation can be induced at specific times is needed. Here, we demonstrate the capability of tRNA synthetase inhibitors, indolmycin and AN3365, to model persistence independently from the use of IFNγ. These tools will also allow comparisons between amino acid stress responses in this unique bacterium.

## Introduction

Chlamydial diseases are significant causes of morbidity in humans. *Chlamydia trachomatis* is the leading cause of bacterial sexually transmitted infections in the world. In 2017, the U.S. Centers for Disease Control and Prevention received over 1.7 million reports of chlamydial infections (1). This number is likely an underestimate due to most infections being asymptomatic and, therefore, undetected (2). The strains responsible for these infections are primarily confined to the urogenital serovars, D-K, but can also contain those of the invasive serovars, L1-L3. Untreated *C. trachomatis* urogenital infections can ascend the genital tract, potentially leading to pelvic inflammatory disease and tubal factor infertility (3). *Chlamydia pneumoniae* is a respiratory pathogen responsible for approximately 10% of community acquired cases of pneumonia. The presence of antibodies in over 50% of adults in the United States, as well as several other countries, suggests infection with *C. pneumoniae* is relatively common (see (4) for extended review). Additionally, long term sequelae such as atherosclerosis and adult-onset asthma have been associated with *C. pneumoniae* infection (5, 6).

Chlamydiae are obligate intracellular bacteria that require a host cell to complete their developmental cycle. Chlamydial development involves interconversion between two distinct developmental forms: the elementary body (EB) and the reticulate body (RB). EBs are infectious, metabolically quiescent, environmentally stable, and compact in size (0.3 µm). RBs are the non-infectious, metabolically active, replicative form that measure approximately 0.8 µm in diameter (as reviewed in (7)). After initial attachment, EBs are internalized into an endocytic vesicle of the host cell and begin primary differentiation into RBs. Soon thereafter, chlamydial proteins are secreted into the vesicle membrane and host cell cytosol to prevent targeting of the chlamydial-containing vacuole to the lysosome. This modified endosome is known as the chlamydial inclusion and is a protective vacuole that masks the invading organisms from host cell defenses for the entirety of their development (8). Following the establishment of the inclusion and primary differentiation into an RB, the organism rapidly multiplies by a polarized budding mechanism (9). *Chlamydia* asynchronously undergo secondary differentiation into EBs until the organisms are released from the cell through lysis or extrusion. The duration of this developmental process is approximately 48 hours for *C. trachomatis* or 72 hours for the slower growing *C. pneumoniae.*

During infection, host immune cells respond to *Chlamydia* by releasing the cytokine interferon-gamma (IFNγ) (10). IFNγ will bind its receptor and activate multiple signaling pathways. The major IFNγ-induced antichlamydial effector in human cells is indoleamine 2,3-dioxygenase (IDO) (11). IDO will catabolize cytosolic tryptophan (trp) into N’-formylkynurenine, a metabolite that cannot be used by *C. trachomatis* or *C. pneumoniae* (11-14). Although IFNγ regulates over 200 human genes (15), IDO expression, with the resulting depletion of available trp (16, 17) and decrease in translation (18), is the driving factor for inhibiting chlamydial growth (Figure 1). This is supported by the ability to restore growth in cell culture by adding additional trp to the media, by pharmacologically inhibiting IDO in the presence of IFNγ, or by using IDO mutant cells (19-21).

**Figure 1.**
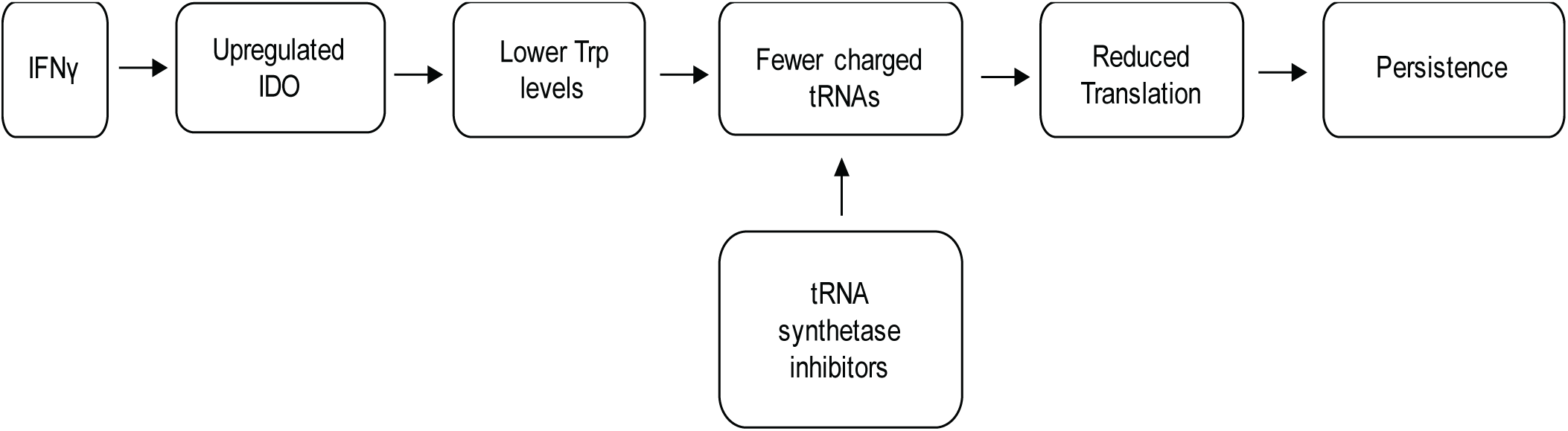
A flowchart illustrating the events leading to IFNγ-mediated persistence. By using tRNA synthetase inhibitors to affect translation, a more direct route to persistence is achieved.

Because *C. trachomatis* and *C. pneumoniae* are trp auxotrophs and depend on host trp to grow, they must respond to this starvation condition to maintain viability (22). Interestingly, *Chlamydia* has eliminated the stringent response (*relA/spoT*), which is used by most eubacteria to respond to amino acid starvation (22, 23). This raises the intriguing question of how they respond to amino acid limitation. Phenotypically, chlamydial RBs transition into an alternative developmental state termed persistence to cope with this stress (16). Persistent chlamydiae are thought to be associated with the chronic sequelae linked to chlamydial diseases. Importantly, various stressors can trigger persistence and not all persistent transcriptomic and proteomic responses are the same (as reviewed in (24)). Nevertheless, persistence models are characterized by bacteria (i) remaining viable yet non-infectious, (ii) being non-replicative, (iii) exhibiting an aberrant morphology, (iv) and reactivating to resume the development cycle after the stress is removed (18, 25). Importantly, and as it relates to IFNγ-induced persistence, transcriptomic and proteomic changes also occur, and these changes are predictable based on the trp content of the transcript or protein (26).

The mechanisms for entry into, maintenance of, and exit from persistence are not known, and investigation of these mechanisms is made difficult by the nature of inducing persistence through IFNγ exposure. Therefore, it is important to be able to study *Chlamydia*’s transition into persistence while simultaneously controlling for as many variables as possible, demonstrating a need for an amino acid starvation model without the confounding variables that come with using IFNγ. In an effort to reproduce and model IFNγ-induced persistence in the absence of IFNγ, the most straightforward approach is to remove trp from the culture medium. However, this will induce host cell autophagy and lysosomal degradation of proteins, which can regenerate amino acid pools that *Chlamydia* can scavenge (27).

We hypothesized that bacterial tRNA synthetase inhibitors would offer an alternative pathway to decrease translation in *Chlamydia* in an amino acid-dependent manner (Figure 1). These inhibitors would, therefore, afford an opportunity to specifically mimic starvation for different amino acids to compare and contrast amino acid starvation responses in *Chlamydia*. We show that combining trp depleted media with the trp analog, indolmycin, is sufficient to induce persistence in *Chlamydia* in cell culture. Indolmycin is a tryptophanyl-tRNA synthetase inhibitor that acts through competitive inhibition as a trp analog (28). Additionally, AN3365, a leucyl-tRNA synthetase inhibitor, was used in this study to investigate the possibility of inducing persistence through the starvation of an amino acid other than trp. AN3365 is an antibiotic in the aminomethyl benzoxaborole class shown to be active against Gram-negative bacteria (29). Unlike indolmycin, AN3365 is a non-competitive inhibitor of bacterial leucyl-tRNA synthetases that locks the protein in its editing conformation, preventing release of charged leucyl-tRNAs (29). These tools will facilitate the modeling of specific amino acid starvation responses in *Chlamydia* without affecting the host cells. Here, we validate the use of these compounds, indolmycin and AN3365, as tools that can be used within the chlamydial field to investigate mechanisms engaged by *Chlamydia* to enter, maintain, and exit persistence in response to amino acid starvation.

## Results

### The bacterial tRNA synthetase inhibitors indolmycin and AN3365 block chlamydial growth

To determine whether the bacterial tRNA synthetase inhibitors indolmycin and AN3365 were effective against *Chlamydia*, we measured inclusion forming units (IFU) as a metric for chlamydial growth in the presence and absence of the inhibitors. Initial empirical experiments with indolmycin, a competitive inhibitor for tryptophanyl-tRNA charging, revealed that it had an impact on growth at 120 µM or higher and in the presence of 1 mg/L trp or lower (data not shown). We chose to perform this and all subsequent experiments with indolmycin by adding it at 120 µM concentration in the absence of trp at 10 hpi. Under these conditions, indolmycin treatment reduced the generation of IFUs to the limit of detection of the assay (Figure 2A). To determine the effective concentration for AN3365, we performed a dose response assay and observed that concentrations in excess of 250 ng/mL (added at 10 hpi) were sufficient to reduce chlamydial growth to basal levels (i.e. near limit of detection, Fig. 2B). Importantly, as a non-competitive inhibitor of leucyl-tRNA charging, AN3365 was effective in the presence of normal media levels of leucine (105 mg/L). AN3365 was also effective at blocking chlamydial growth when added at various time points up to 12 hpi during the developmental cycle (Fig. 2C). From these data, we conclude that indolmycin and AN3365 effectively blocked chlamydial growth. Nevertheless, a lack of recoverable IFUs suggests one of three outcomes: i) complete loss of viability, ii) decrease in the rate of development (i.e. prolonged RB-only phase), or iii) entry into persistence.

**Figure 2.**
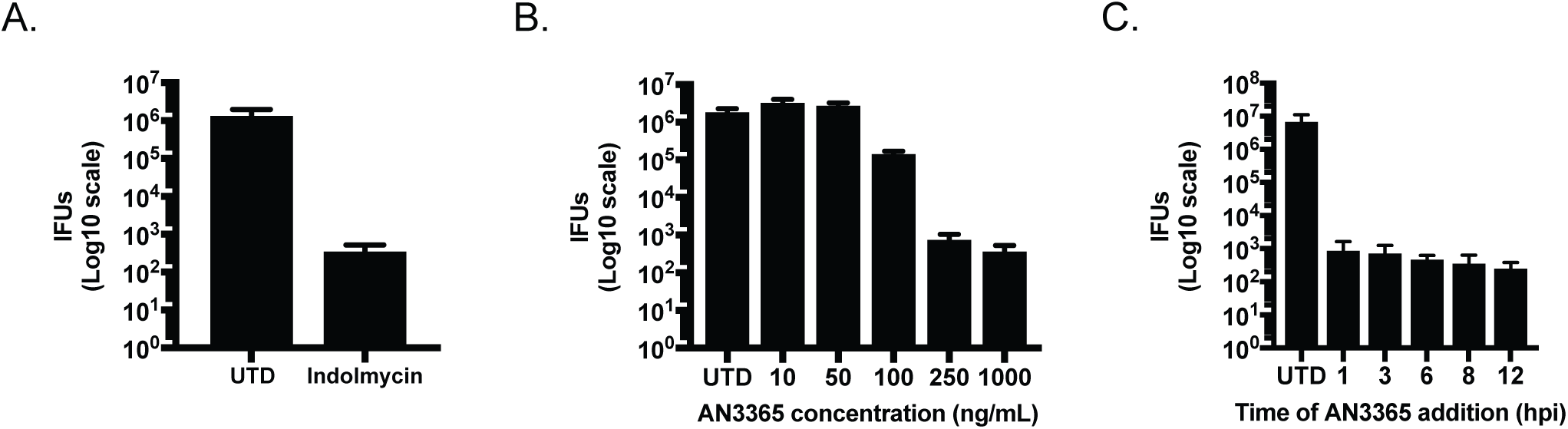
Indolmycin and AN3365 effectively reduce recoverable infectious elementary bodies. For all experiments, HEp-2 cells were infected with *C. trachomatis* L2, and inclusion forming units (IFUs) were collected at 24 hpi and titrated on a fresh monolayer of HEp-2 cells in the absence of antibiotics. A) Indolmycin was added at 120 µM at 10 hpi. B) Effect of various doses of AN3365 on IFU production when added at 10 hpi. C) AN3365 was added at different times post infection at 1 µg mL^-1^ as indicated. Data are representative of three separate biological replicates. Error bars represent the standard deviation between biological replicates.

### Indolmycin and AN3365 induce morphological aberrance in *C. trachomatis*

To help distinguish between the possible reasons for a loss in recoverable IFUs, immunofluorescence microscopy was used to analyze chlamydial morphology (Fig. 3). HEp-2 cells were infected with *C. trachomatis* serovar L2, treated or not with the indicated compounds at 10 hpi or pre-treated with IFNγ as described in Materials and Methods, and fixed and processed for imaging at 24 hpi. Both indolmycin and AN3365 treatment resulted in noticeably larger individual organisms, similar to that observed during IFNγ treatment (16). Electron microscopic analysis of organisms treated with inhibitors also revealed aberrant morphology (Suppl. Fig. 1). Interestingly, the labeling of MOMP was not uniform around the membrane of the organisms in the treated cells. The *ompA* gene encodes 30 leu (L) and 7 trp (W) codons. Decreased availability of L or W may therefore negatively impact translation of MOMP that in turn may lead to non-uniform localization along the organism’s membrane (see also (16)). Conversely, Hsp60_1, which contains 0 W codons, appeared abundant in both indolmycin and IFNγ-treated samples. Interestingly, Hsp60_1 is also abundant in the AN3365 treated sample despite the presence of 45 L codons. These observations are consistent with what has been previously described during IFNγ-induced persistence (12). Taken together, these morphological data in conjunction with the IFU data presented above support the conclusion that the organisms are non-replicative, display an aberrant morphology, and are not proceeding through the normal developmental cycle.

**Figure 3.**
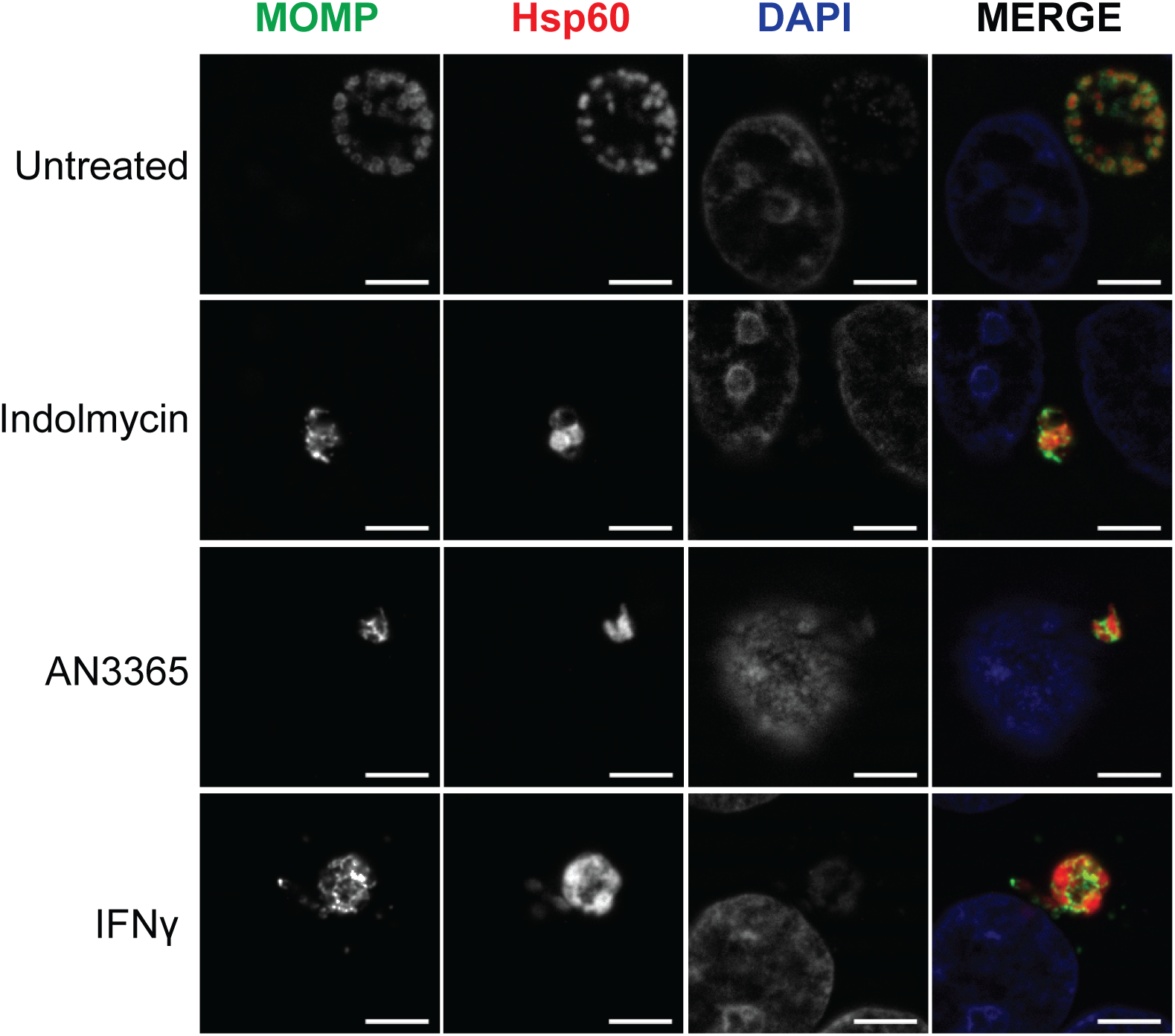
Inhibition of tRNA synthetase results in morphological aberrance in *C. trachomatis* L2. Representative images of HEp-2 cells infected with *C. trachomatis* and treated or not as indicated. Cells were fixed and stained 24 hpi using primary antibodies to Major Outer Membrane Protein (MOMP) and chlamydial Hsp60_1. Indolmycin and AN3365 treatments resulted in smaller inclusions and morphologically aberrant organisms similar to IFNγ treated organisms. All images were acquired on Zeiss LSM 800 confocal microscope with Airyscan at 63x optical magnification with 2x digital zoom. Scale bars represent 5 µm.

### Removal of indolmycin and AN3365 rescues *C. trachomatis* growth and morphology

A key characteristic of persistence is its reversibility such that the organism can revert back to a developmentally competent RB. To determine whether or not *C. trachomatis* remains viable under indolmycin or AN3365 treatment, we attempted to rescue the organisms by removing each treatment (Fig. 4). HEp-2 cells were infected with *C. trachomatis* serovar L2 and treated or not with the designated compounds at 10 hpi. Treatment media was aspirated at 24 hpi, and samples were washed 3x with DPBS before replenishing with DMEM. After 24 and 48 hour recovery periods following drug removal, IFUs were quantified and organism morphology was assessed. In the presence of indolmycin and AN3365, chlamydial growth was inhibited up to 72 hpi (Figure 4A). After removal of these compounds from the medium, chlamydial growth was restored as represented by a logarithmic increase in IFUs at 48 (24h post recovery) and 72 (48h post recovery) hpi. These IFU data were corroborated by the detection of normal morphological forms at these time points that were indistinguishable from 24 and 48 hpi untreated organisms (Figs. 4B-D). Moreover, recoverable IFUs were intermediate between that observed at the 24 and 48 hpi untreated groups. This is expected when considering treatment time at 10 hpi essentially pauses the development cycle. After 24 hours of recovery, the organisms have undergone approximately 34 hours of development. We conclude that the effects of indolmycin and AN3365 are reversible, consistent with what is observed during the removal of IFNγ from persistent cultures (18, 25, 30). Collectively, the data presented in Figures 2-4 indicate that the tRNA synthetase inhibitors induce a persistent growth state that is reversible upon removal of the compounds.

**Figure 4.**
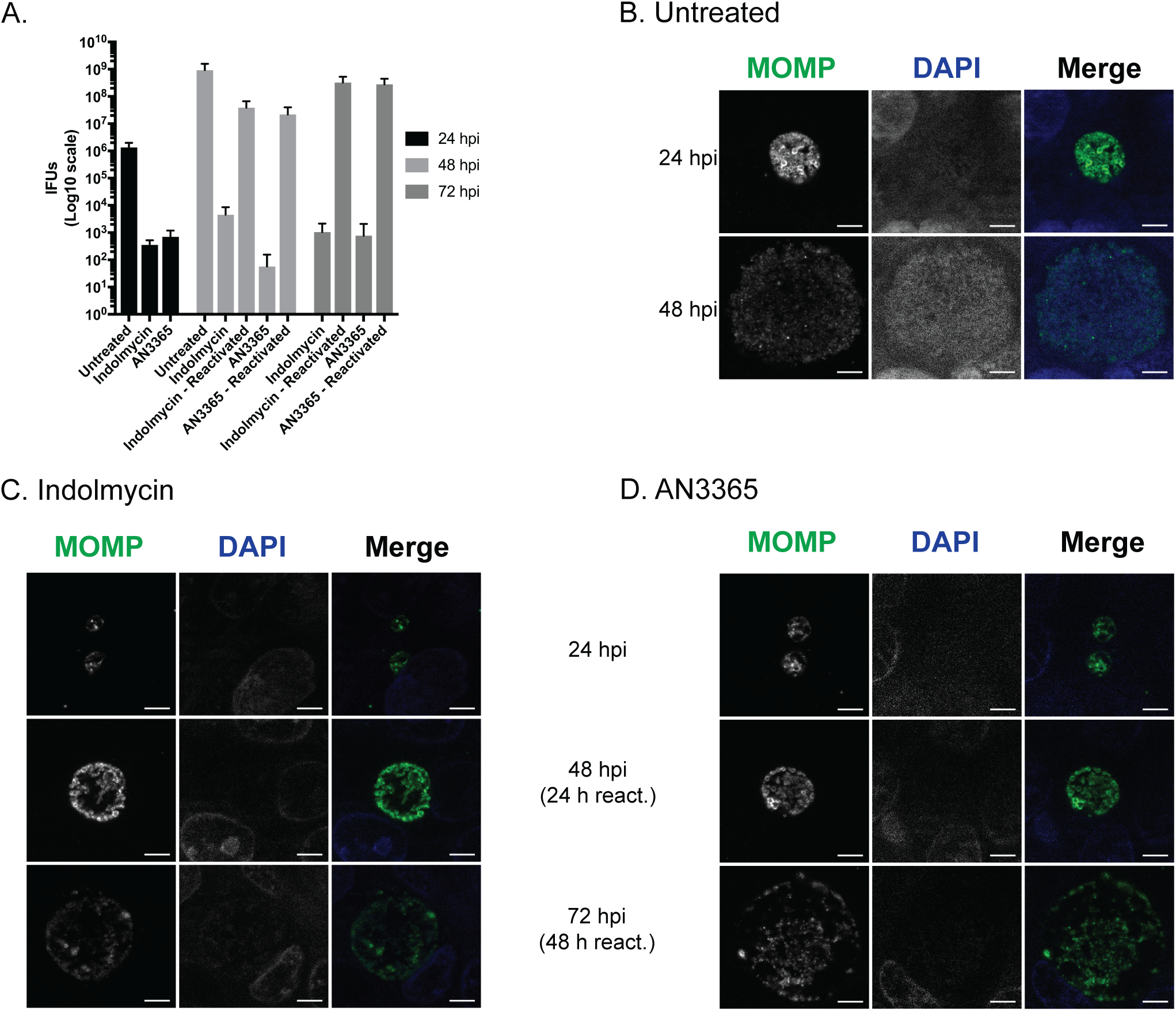
Removal of indolmycin and AN3365 allows reactivation from persistence. HEp-2 cells were infected with *C. trachomatis* and treated or not with the designated tRNA synthetase inhibitor at 10 hpi. DMEM was removed at 24 hpi followed by three DPBS washes and replenishment with unmodified DMEM. Cultures were allowed to recover for an additional 24 or 48 hours before fixation. A) IFU samples were collected from replicate wells at designated points throughout the experiment. Error bars represent variability between three biological replicates. B) Representative images of untreated *C. trachomatis* infected HEp-2 cells at 24 or 48 hpi. C & D) Representative images of indolmycin (C) or AN3365 (D) treated *C. trachomatis* infected HEp-2 cells at 24, 48 (24h reactivation = react.), or 72 (48h react.) hpi. All images were acquired on an AXIO Imager.Z2 with ApoTome.2 at 100x magnification. Scale bars represent 5 µm.

### Indolmycin and AN3365 induce morphological aberrance in *C. pneumoniae*

Although the underlying mechanisms of persistence are unknown, we hypothesize that they are conserved between *C. trachomatis* and *C. pneumoniae. C. pneumoniae* is a slower growing species compared to *C. trachomatis*, particularly the L2 serovar, suggesting it would be equally, if not more, sensitive to tRNA synthetase inhibitors. Therefore, we suspected the use of indolmycin and AN3365 on *C. pneumoniae* would produce a phenotype similar to that observed during IFNγ exposure. To test this, we infected HEp-2 cells with *C. pneumoniae* AR39. Samples were treated or not with the designated compound at 24 hpi or with IFNγ at time of infection. As seen in Figure 5, the morphologies of organisms treated with indolmycin or AN3365 closely resemble those treated with IFNγ. We conclude from these data that tRNA synthetase inhibitors are broadly applicable to study persistence in *Chlamydia* species.

**Figure 5.**
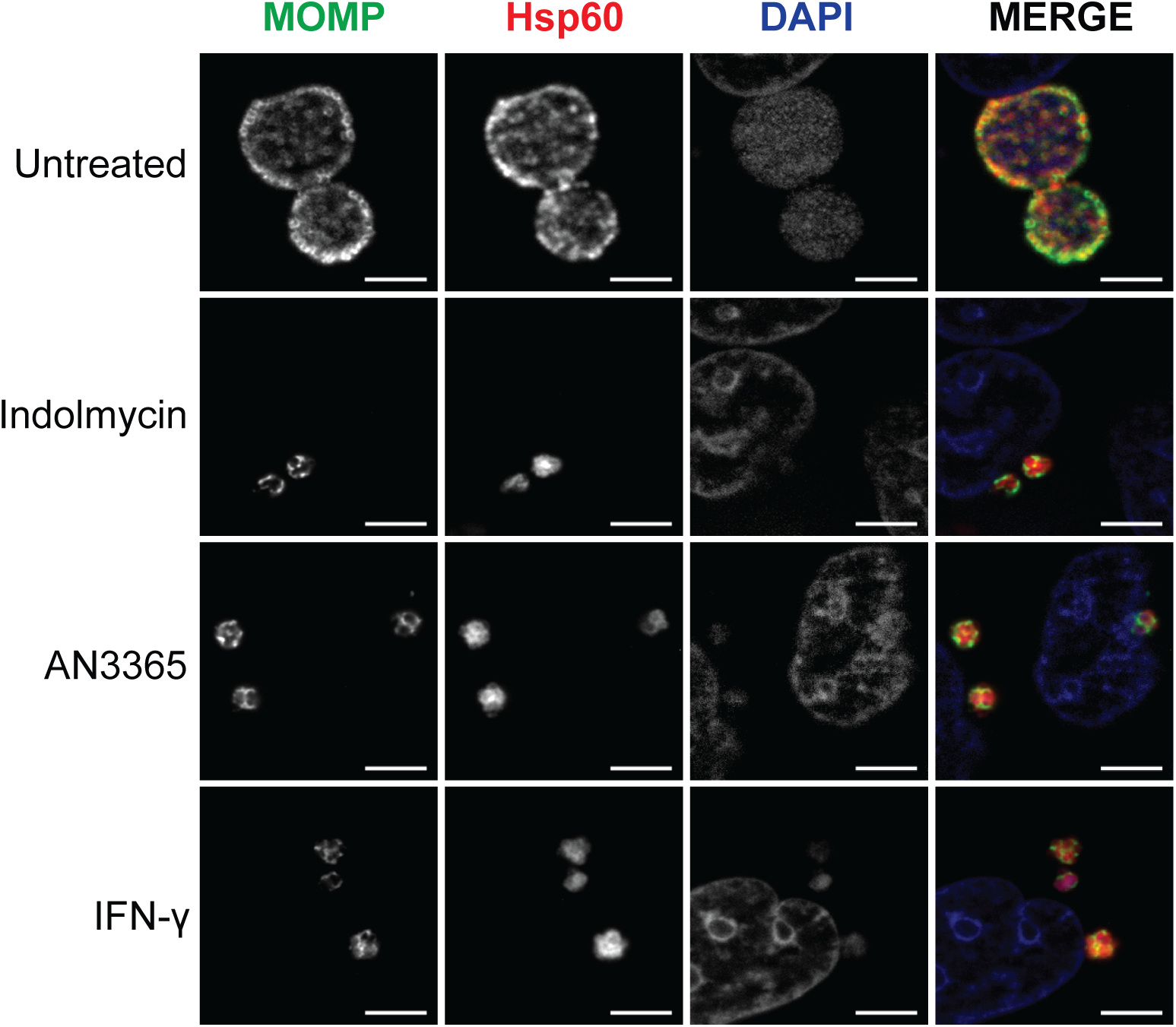
Immunofluorescent images of HEp-2 cultures infected with *C. pneumoniae* at 48 hpi show morphological similarity between indolmycin and AN3365 treatments with IFNγ induced persistent organisms. Indolmycin and AN3365 were added at 24 hpi. IFNγ was added at time of infection. All images were acquired on an AXIO Imager.Z2 with ApoTome.2 at 100x magnification. Scale bars represent 5 µm.

### The bacterial tRNA synthetase inhibitors induce transcriptional changes consistent with IFNγ-mediated persistence

To determine whether the tRNA synthetase inhibitors induce transcriptional changes consistent with persistence, nucleic acids were isolated from infected cultures, and the abundance of selected transcripts was measured and normalized to genomic DNA content. Increased transcription of *euo* has been previously associated with IFNγ-mediated persistence (18, 30). This elevated transcriptional response was observed in both *C. trachomatis* (beginning at 4h post treatment) and *C. pneumoniae* when exposed to either indolmycin or AN3365 treatments and resembled IFNγ-induced persistence (Fig. 6A). In addition to *euo, groEL_1* has also been implicated in persistence and general stress response (30). We analyzed the abundance of *groEL_1* transcripts in both *C. trachomatis* and *C. pneumoniae* under each treatment (Fig. 6B). In agreement with previous findings, *groEL_1* transcripts remained unchanged (<1.5x difference) in *C. trachomatis* between 10 and 24 hpi in IFNγ-treated samples, whereas *groEL_1* transcripts decreased in abundance during the same timeframe in the untreated samples (18). Interestingly, indolmycin-treated samples closely mirrored the effects of IFNγ while AN3365 treatment resulted in a 4-5 fold increase of *groEL_1* transcripts at the 24 hour time point. Conversely, for *C. pneumoniae*, all treatment conditions followed a similar transcript pattern as in untreated samples for *groEL_1*. This is also consistent with previous observations for IFNγ-induced persistence in this organism (18). Additionally, indolmycin treatment resulted in a rapid (4h) and large increase in *trpB* transcripts (*C. pneumoniae* does not encode *trpB*), a gene that is repressed under trp-replete conditions (Fig. 6C and (31)). This suggests *C. trachomatis* has detected and is quickly responding to a lack of trp. Moreover, there is a noticeable increase in *trpB* transcripts at 24 hpi following AN3365 treatment (3-4 fold increase). Considering the lack of change observed at 14 hpi, the increase of *trpB* transcripts at 24 hpi may be an indirect effect caused by a cascade of responses rather than an immediate reaction to leu limitation. It is important to note that while *C. trachomatis* detects trp limitations when cultured in trp-deplete media (no indolmycin), the abundance of *euo* transcripts does not change (Suppl. Fig. 2). Therefore, removing trp from the medium cannot be used as a reliable IFNγ-free model for persistence, particularly for faster growing species and strains of *Chlamydia*.

**Figure 6.**
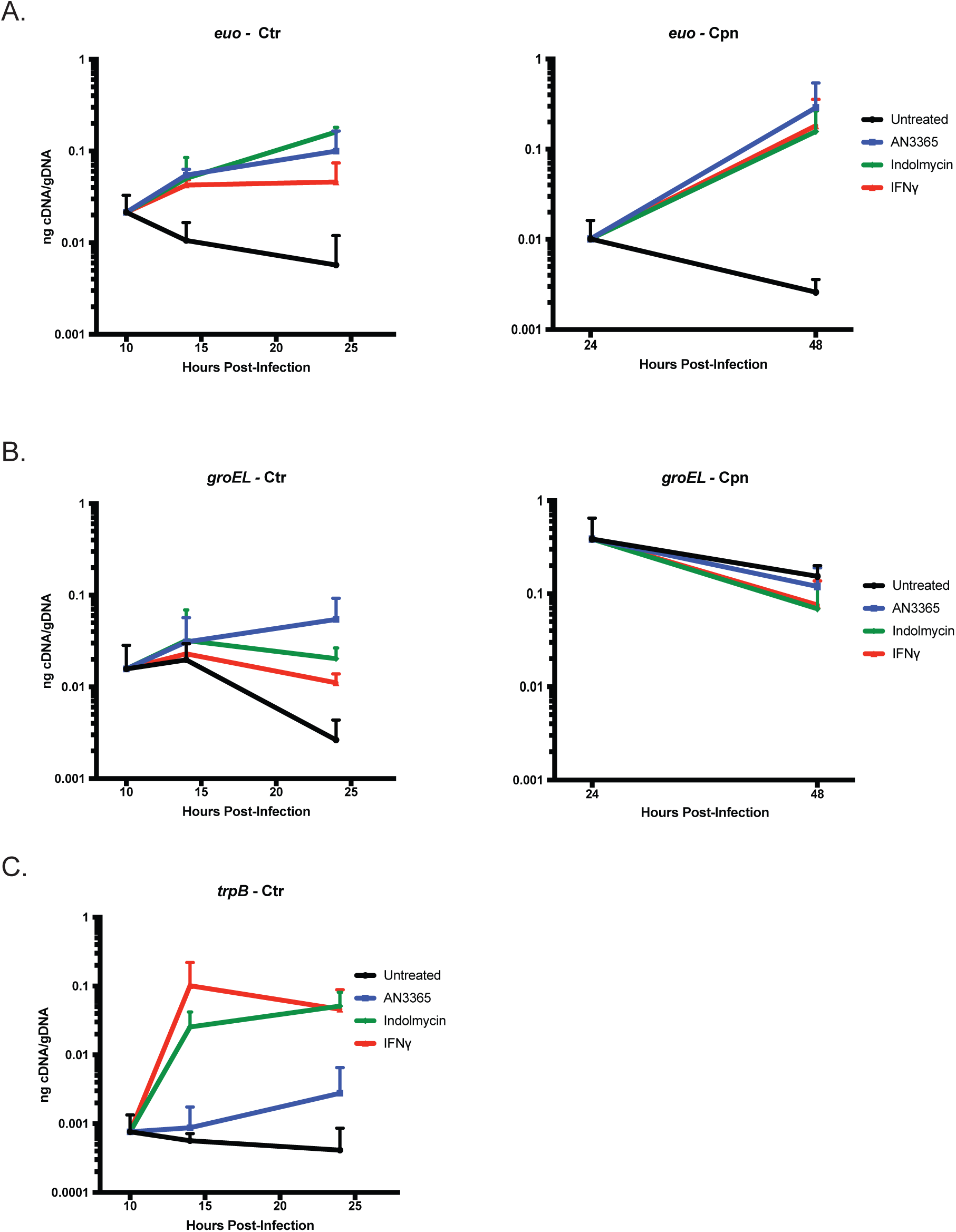
Transcriptional changes in Chlamydia consistent with persistence can be detected during indolmycin and AN3365 treatment. A) Transcripts of euo are elevated in C. trachomatis (Ctr) or C. pneumoniae (Cpn) following treatments with indolmycin, AN3365, or IFNγ. B) Under standard conditions, groEL_1 transcripts in C. trachomatis decrease between 10 and 24 hpi. However, treatment with indolmycin results in unchanged transcript levels, similar to what is seen in IFNγ treated samples, while AN3365 treatment results in higher transcript levels. In C. pneumoniae, no significant change is seen between 24 and 48 hours under any treatment, in agreement with previous reports investigating IFNγ exposure. C) trpB transcripts accumulate at 14 hpi as expected in indolmycin and IFNγ treated samples, but not in AN3365. At 24 hpi, AN3365 treated samples exhibit a slight (3-4 fold) increase in trpB.

### Transcript levels of the 3’ end of the *ytg* operon are reduced in *Chlamydia* during indolmycin or AN3365 treatment

To determine whether indolmycin and AN3365 could mimic a more nuanced characteristic of IFNγ-induced persistence, we looked to the *ytgABCD* operon (26). As previously described by Ouellette et al. (32), IFNγ treatment results in Rho-dependent polarization of the *ytg* operon. This results in a skewed ratio of *ytgA:ytgD* transcripts. During the normal developmental cycle, this ratio is approximately 5-10 but increases to over 30-fold or higher during IFNγ-mediated trp starvation. We hypothesized that this skew was caused by ribosome stalling along the transcript due to the lack of charged tRNAs, which allows Rho to bind internal *rut* sites to terminate transcription prematurely (32). Of note, within the *ytgC* gene is the presence of three tandem W codons, and the gene encodes additional W residues. L residues are highly abundant in the operon with 98 total residues in *ytgB* and *ytgC* and a total of 11 LL motifs in *C. trachomatis*. We reasoned that both indolmycin and AN3365 should produce the same phenotype as a result of the stalling of ribosomes on trp codons, in the case of indolmycin treatment, or leu codons, in the case of AN3365 treatment, respectively. We quantified *ytgA* and *ytgD* transcripts at different times after addition of the tRNA synthetase inhibitors (at 10hpi for Ctr, 24hpi for Cpn) and compared the ratios to those from IFNγ-treated cultures. As anticipated, both indolmycin and AN3365 treated samples resembled the IFNγ-induced persistent state with regards to the polarity of the *ytg* operon, shown in Figure 7 as a disproportionate level of *ytgA* to *ytgD* transcripts. Interestingly, differences in the Ctr *ytgA:ytgD* ratio were observed within four hours of treatment (14hpi) and were more pronounced for L limitation (AN3365) than for W limitation (indolmycin or IFNγ) (Fig. 6A). This is consistent with the larger number of L versus W residues in the operon. However, by 24hpi, the *ytgA:ytgD* ratio for all treatments was similar, possibly reflecting a recovery in read-through potential in AN3365-treated cultures. For Cpn, all treatments resulted in the expected increase in *ytgA* transcripts in proportion to *ytgD* transcripts when measured at 48hpi (after treatment at 24hpi). Overall, these data demonstrate that the tRNA synthetase inhibitors recapitulate key characteristics of amino acid starvation in *Chlamydia*.

**Figure 7.**
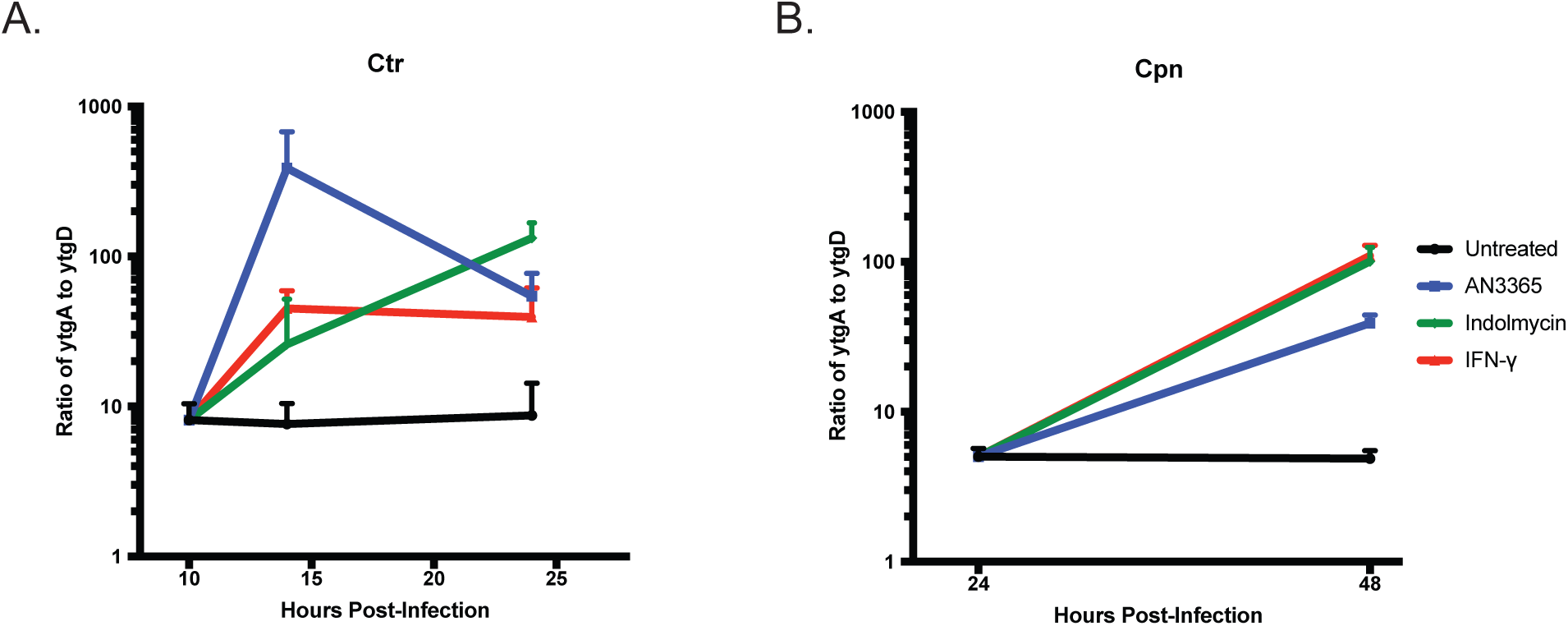
Transcriptional analysis shows a decrease in readthrough efficiency of the *ytgABCD* operon during indolmycin, AN3365, or IFNγ treatment in A) *C. trachomatis* (Ctr) or B) *C. pneumoniae* (Cpn). RT-qPCR was performed to determine ng of cDNA of both *ytgA* and *ytgD.* Each was normalized to gDNA collected from replicate wells and expressed as a ratio of ng cDNA per ng gDNA of *ytgA* over *ytgD.*

## Discussion

Persistence is an important but poorly understood aspect of chlamydial diseases. The immunological basis for inducing persistence in cell culture was first described by Beatty et al. in 1993, who described the effects of IFNγ, and its reversibility, on chlamydial growth and morphology in human cell lines (12, 16, 25). These effects were connected with the ability of human IFNγ to induce a tryptophan-limiting environment in the cell by activating IDO expression (12). Broadly speaking, these effects are likely mediated by the inability to efficiently translate key proteins enriched in trp residues (see Fig.1 and (18, 33, 34)). More recent studies have characterized transcriptional and translational changes associated with IFNγ-mediated persistence (18, 30). More importantly, these “persistence” characteristics as defined in cell culture models have recently been observed in patient samples (13). This underscores the need to have a better mechanistic understanding of how amino acid limitation results in a persistent phenotype.

In 2011, the Clarke group published the first study demonstrating stable transformation of *Chlamydia trachomatis* serovar L2 (35). This advance, common for decades in other bacterial systems, has allowed fluorescent tagging of target proteins and reverse genetic tools to be applied to *Chlamydia* (36-40). However, serovar L2 is among the fastest growing strains of *C. trachomatis* (41), and slower growing strains and species of *Chlamydia* have proven more difficult to transform. As it relates to genetic studies of IFNγ-induced persistence, this creates a hurdle. When IFNγ is added to cell cultures, removal of trp from the cytosol by IDO is gradual and takes approximately 24 h. This is a time during the L2 developmental cycle when RBs are differentiating to EBs and EBs are rapidly accumulating (e.g. Fig. 2A). For IFNγ to be effective, pretreatment of host cells with IFNγ prior to infection is required, yet this strategy is inconsistent. To induce a persistent state in serovar L2, we recently published a protocol that described the pretreatment of cells with IFNγ prior to infection followed by the addition of IFNγ-conditioned medium at 10 hpi (32). This allows IDO to be sufficiently expressed and trp to be catabolized, resulting in small inclusions that contain relatively few aberrant organisms approximately 24 hours post infection (hpi). While this protocol elicited reproducibly persistent forms for us, differences between labs, the cell types used, and batches of IFNγ (which require careful titration for effective dose) may not make it easily transferable to other systems. This can lead to discrepancies in findings since too much IFNγ exposure prevents primary differentiation of chlamydial EBs to RBs, while too little results in mixed populations of persistent and normal organisms within a culture (20). In sum, the field would benefit greatly from a tool that allows reproducible induction of persistence and that minimizes confounding variables while maximizing control and flexibility of experimental design parameters.

Here, we sought to evaluate the effects of characterized bacterial tRNA synthetase inhibitors (28, 29, 42) for their ability to induce persistence in *Chlamydia* as a first step in developing systems that would allow us to mechanistically address this alternative growth state. Such an approach would offer immediate advantages over IFNγ-mediated tryptophan limitation in that a translation block (i.e. starvation mimicking condition) could be induced immediately upon addition of the inhibitors. By using *E. coli* tRNA turnover rates as a guide, we hypothesize that the pool of charged trp-tRNA or leu-tRNA would be depleted within seconds after treatment (43). This would allow more direct comparisons between research groups with less variability in experimental systems. In particular, the use of inhibitors circumvents the host cell’s ability to regenerate amino acid pools through autophagy, which would occur in conditions where an amino acid is omitted from culture medium. Indeed, we observed that, under such conditions, although the absence of trp was sensed, as demonstrated by increased *trpB* transcripts, *euo* transcripts, a marker of persistence, did not increase (Suppl. Fig. 2). This suggests that, for serovar L2, simply omitting amino acids from the culture medium is not sufficient to induce a *bona fide* persistence response.

These data indicate the ability of indolmycin and AN3365 to induce persistence by limiting *Chlamydia*’s use of a single specific amino acid by blocking the charging of its cognate tRNA. Considering the parallels in morphology, transcriptional response, and mode of stress caused by these compounds in comparison to IFNγ, we conclude that the use of indolmycin and AN3365 in place of IFNγ is a viable method to study amino acid starvation stress responses. Interestingly, indolmycin treatment replicates the key transcriptional and morphological phenotypes associated with IFNγ-induced persistence, further supporting that tryptophan limitation is the main antichlamydial inhibitory mechanism of IFNγ in human cells (11, 19). The transcriptional changes of *C. pneumoniae* in response to the inhibitors more closely mirrored IFNγ-mediated persistence than *C. trachomatis* did. Given the slower growth rate of *C. pneumoniae*, this is not surprising. The greater heterogeneity in transcription responses between indolmycin and IFNγ in *C. trachomatis* may be due to the organism’s quicker developmental cycle and asynchronous nature. That being said, the difference between the tryptophan starvation condition in *C. pneumoniae* versus *C. trachomatis* was not more than four-fold and showed the same trends overall.

AN3365-induced leu starvation displayed noteworthy differences from the trp starvation conditions. Firstly, *groEL_1* transcripts generally increased during the analysis in *C. trachomatis*. This is consistent with what we previously characterized as codon-dependent transcriptional changes during amino acid starvation as Hsp60_1 contains 45 L residues (26). Secondly, *trpB* transcripts were not increased 4h after treatment, as expected since leu starvation should not activate expression of the *trpRBA* operon. However, 14h post treatment, there was an approximately 3-fold increase in *trpB* levels, suggesting some de-repression of the operon. TrpR contains 11 leu residues including one LL motif. Therefore, the inability to efficiently translate the repressor over time may allow for gradual transcriptional activation of the operon. Alternatively, the recently described role for the iron-sensitive repressor, YtgR, in blocking transcription of *trpBA*, may also be important (44). The YtgCR protein contains approximately 60 leu residues with multiple LL motifs that likely prevent efficient translation of this sequence. This is under investigation, but we also observed transcriptional changes in the *ytg* operon (see below).

We recently demonstrated differences in transcript levels between the 5’ and 3’ ends of large monocistronic and polycistronic transcripts (26). We further connected this to Rho-dependent polarity prematurely terminating transcription in trp-codon rich transcripts during IFNγ-mediated trp starvation (32). To determine whether the tRNA synthetase inhibitors could replicate the destabilization of the 3’ end of a large transcript, we analyzed the abundance of the *ytgA* and *ytgD* transcripts. Consistent with what was previously observed for IFNγ-mediated trp starvation, both indolmycin and AN3365 caused a disparity in transcript abundance between the 5’ and 3’ ends of the *ytg* operon. Interestingly, AN3365 caused a quick destabilization of *ytgD* transcripts in *C. trachomatis* before recovering to levels observed in trp starvation conditions. The reasons for this are not clear but are under investigation.

The tools described here to mimic specific amino acid starvation states, by blocking tRNA charging, in the absence of chemokines or other significant alterations to culture conditions will facilitate broad comparisons of chlamydial persistence between species. For example, *C. caviae* resists IFNγ-mediated persistence by recycling the product of trp degradation, N-formylkynurenine, through a trp scavenging pathway (45). Indolmycin treatment will facilitate studies of trp starvation responses in this species. Likewise, these treatments can be used in mouse cells, where IDO is not the primary antichlamydial effector (10, 46). Also of interest are other intracellular pathogens such as *Coxiella* or *Rickettsia*, as studying their response mechanisms to amino acid starvation could lead to a greater understanding of evolutionary strategies employed by obligate intracellular pathogens, which typically lack functional stringent responses, to adapt to this stress.

Following the validation of these compounds as tools to study persistence, we aim to further investigate the role of amino acid limitation in regulating the persistent state. With the ability to limit an amino acid other than trp, we can more rigorously test the hypothesis that trp limitation increases transcription of trp codon containing genes to determine if this is a broad response to amino acid limitation or perhaps something specific to trp (26). We can now also apply genetic tools to study amino acid starvation responses in *C. trachomatis* L2.

## Materials and Methods

### Organisms and cell culture

The human epithelial cell line HEp-2 was routinely cultivated at 37°C with 5% CO_2_ in Dulbecco’s modified Eagle medium (DMEM; Gibco, Dun Laoghaire, Ireland) supplemented with 10% FBS. The HEp-2 cells were a kind gift of H. Caldwell (NIH/NIAID). *C. trachomatis* serovar L2 and *C. pneumoniae* AR39 EBs were harvested from infected HEp-2 cell cultures at 37°C and 35°C, respectively, with 5% CO_2_ and density gradient purified. Purified EBs were titered for infectivity by determining inclusion-forming units (IFU) on fresh cell monolayers. All bacterial and eukaryotic cell stocks were confirmed to be *Mycoplasma* negative using the LookOut Mycoplasma PCR Detection Kit (Sigma, St. Louis, MO).

Indolmycin was purchased from Cayman Chemical (Ann Arbor, MI) and resuspended to 120 mM in dimethyl sulfoxide (DMSO; Sigma). Aliquots were kept at −80°C and used only once to avoid freeze-thawing. Indolmycin was used at 120 µM and added at 10 hpi (*C. trachomatis*) or 24 hpi (*C. pneumoniae*) in all experiments. Immediately prior to treatment, cell medium was replaced with DMEM lacking trp (made in-house) to enhance the inhibitory effects of indolmycin. DMEM lacking trp was made using 10% fetal bovine serum that had been dialyzed to remove any additional amino acids. All custom medium components were purchased from Sigma.

AN3365 was purchased from Cayman Chemical and resuspended to 5 mg mL^-1^ in DMSO. Aliquots were kept at −20°C and allowed one additional freeze-thaw. AN3365 concentration was titrated to induce persistence without completely stalling development and was used at 1 µg mL^-1^ with treatment at 10 hpi (*C. trachomatis*) or 24 hpi (*C. pneumoniae*). No modifications to DMEM were necessary. In some experiments with *C. trachomatis*, AN3365 was added at different concentrations or at different times post-infection as indicated, with IFU samples collected at 24hpi.

Recombinant human interferon gamma (IFNγ) was purchased from Cell Sciences (Canton, MA) and resuspended to 100 µg ml^-1^ in 0.1% bovine serum albumin (BSA; Sigma) diluted in water. Aliquots were frozen at −80°C and used only once to avoid freeze-thawing. IFNγ was titrated for its effect to induce persistence without killing the bacteria and, in our experiments, 0.5 ng ml^-1^ was added to cells approximately 11 h prior to infection. Medium was replaced at 10 hpi with IFNγ-conditioned medium (ICM) to induce persistence in *C. trachomatis* as described (32). ICM was prepared by adding 2 ng ml^-1^ IFNγ to uninfected HEp-2 cells for approximately 54 h prior to collection and filtration of the medium. *C. pneumoniae* experiments were treated with 2 ng ml^-1^ at the time of infection.

### Inclusion Forming Unit (IFU) assays

Infectious progeny were determined based on inclusions formed from a secondary infection. Primary infection samples were harvested by scraping cells in 2 sucrose-phosphate (2SP) solution. Samples were lysed via a single freeze-thaw cycle and allowed to infect a fresh cell monolayer. Titers were enumerated by immunofluorescence.

### Immunofluorescent microscopy

Cells were cultured on glass coverslips in 24-well tissue cultures plates and infected with *C. trachomatis* at an MOI of 1 or *C. pneumoniae* at an MOI of 2. All cells were fixed in 100% methanol. Organisms were stained using a primary goat or mouse antibody specific to either *C. trachomatis* or *C. pneumoniae* major outer membrane protein (MOMP), respectively, and a donkey anti-goat or anti-mouse secondary antibody conjugated to Alexa Fluor 488 (Jackson Labs, Bar Harbor, Maine). Where applicable, a primary mouse (Ctr) or rabbit (Cpn) antibody specific to chlamydial Hsp60 was also used in conjunction with a secondary donkey anti-mouse or anti-rabbit antibody conjugated to Alexa Fluor 594 (Jackson Labs).

### Nucleic acid extraction and RT-qPCR

RNA extraction was performed on infected cell monolayers using TRIzol (Invitrogen/ThermoFisher). Samples were treated with Turbo DNAfree (Ambion/Thermofisher) according to the manufacturer’s instructions to remove DNA contamination. cDNA was synthesized from DNA-free RNA using random nonamers (New England BioLabs, Ipswich, MA) and SuperScript III RT (Invitrogen/ThermoFisher) per manufacturer’s instructions. Reaction end products were diluted 10 fold with molecular biology-grade water, aliquoted for later use, and stored at −80°C. Equal volumes of each reaction mixture were used in 25 µL qPCR mixtures with SYBR green master mix (Applied Biosystems) and quantified on a Quant Studio 3 (Applied Biosystems/ThermoFisher) using the standard amplification cycle with a melting curve analysis. Results were compared to a standard curve generated against purified *C. trachomatis* L2 or *C. pneumoniae* genomic DNA as appropriate. DNA samples were collected from replicate wells during the same experiments using the DNeasy Blood and Tissue kit (Qiagen, Hilden, Germany). Equal total DNA quantities were used in qPCR with a *groEL1* primer set to quantify chlamydial genomes. Genome values were used to normalize respective transcript data. RT-qPCR results were normalized for efficiency with typical results demonstrating r^2^>0.995 and efficiencies greater than 90%.

### Reactivation

To determine the possibility of recovery from persistence, samples were treated or not with AN3365 or indolmycin as described above. After 24 hours post infection, all samples were washed three times with DPBS and given fresh medium. Samples were allowed to recover for an additional 24 or 48 hours before collection for IFU assay or fixation for immunofluorescent microscopy.

### Electron Microscopy

HEp-2 cells were cultured in a 6-well plate and infected with *C. trachomatis* serovar L2 at an MOI of 2.5. Samples were treated or not with indolmycin or AN3365 at 10 hpi. Cells were trypsinized at 24 hpi and pelleted at 500 xg for 5 minutes. Pellets were washed 3x using DPBS. Following the final wash, pellets were resuspended in 1 mL of fixative containing 2% glutaraldehyde, 2% paraformaldehyde, and 0.1M Sorenson’s phosphate buffer, pH 7.2. Post fixation was carried out using 1% Osmium Tetroxide followed by a dehydration series in increasing EtOH concentrations. 90 nm sections were cut using a Leica UC6 Ultramicrotome with a Diatome diamond knife. Sections were stained in 2% Uranyl Acetate and Reynold’s Lead Citrate. Images were collected on an FEI Tecnai G2 TEM operated at 80 Kv.

## Acknowledgements

This work was supported by start-up funds from the University of Nebraska Medical Center as well as a CAREER award (1810599) from the National Science Foundation to SPO. We thank Dr. H. Caldwell (NIAID/NIH) for eukaryotic cell stocks and the antibody to *C. pneumoniae* MOMP, Dr. R. Morrison (UAMS) for the antibody to chlamydial Hsp60_1, Dr. E. Rucks (UNMC) for the antibody against *Chlamydia* and for critical review of the manuscript, and Dr. R. Carabeo (UNMC) for critical review of the manuscript. We would also like to thank Tom Bargar and Nicholas Conoan of the Electron Microscopy Core Facility (EMCF) at the University of Nebraska Medical Center for technical assistance. The EMCF is supported by state funds from the Nebraska Research Initiative (NRI) and the University of Nebraska Foundation, and institutionally by the Office of the Vice Chancellor for Research.

This publication’s contents and interpretations are the sole responsibility of the authors.

We declare that we have no conflict of interest.

---

**Supplemental Figure 1.** Electron micrograph images were collected to examine the morphological impact of indolmycin and AN3365 on *C. trachomatis*. HEp-2 cells were infected with *C. trachomatis* at an MOI of 2.5 and treated or not with the denoted tRNA synthetase inhibitor. Samples were collected and fixed at 24 hpi. Images were acquired on an FEI Tecnai G2 TEM operated at 80 Kv. Scale bars represent 2 µm, 500 nm, and 500 nm, respectively.

**Supplemental Figure 2.** Trp deplete media induces an increase in *trpB*, but not *euo*, transcripts. HEp-2 cultures were infected with *C. trachomatis* at an MOI of 1. At 10 hpi, DMEM was replaced with standard DMEM (Untreated), DMEM lacking trp (No trp), or DMEM lacking trp with 120 µM indolmycin (Indolmycin). RNA transcripts were analyzed via RT-qPCR to compare the efficacy of trp deplete media in inducing persistence, as measured by increased *euo* transcript levels.

**Supplemental Table 1.** All primer sequences used for qPCR analysis of a given transcript.

